# A HicA toxin-based counter-selection marker for allelic exchange mutations in Fusobacterium nucleatum

**DOI:** 10.1101/2023.01.20.524997

**Authors:** GC Bibek, Peng Zhou, Chenggang Wu

## Abstract

The study of fusobacterial virulence factors has dramatically benefited from the creation of various genetic tools for DNA manipulation, including the *galK-based* counterselection for in-frame deletion mutagenesis in *Fusobacterium nucleatum* that was recently developed. However, this method requires a host lacking the *galK* gene, which is an inherent limitation. To circumvent this limitation, we explored the possibility of using the *hicA* gene that encodes a toxin consisting of a HicAB toxin-antitoxin module in *Fusobacterium periodonticum* as a new counter-selective marker. Interestingly, the full-length *hicA* gene is not toxic in *F. nucleatum*, but a truncated *hicA* gene version lacking the first six amino acids is functional as a toxin. The toxin expression is driven by an *rpsJ* promoter and is controlled at its translational level using a theophylline-responsive riboswitch unit. As a proof of concept, we created markerless in-frame deletions in the fusobacterial adhesin RadD gene within the *F. nucleatum rad* operon and the *tnaA* gene that encodes the tryptophanase for indole production. After vector integration, plasmid excision after counterselection appeared to have occurred in 100% of colonies grown on theophylline-added plates and resulted in in-frame deletions in 50% of the screened isolates. This *hicA*-based counterselection system provides a robust and reliable counterselection in wild-type background *F. nucleatum* and should also be adapted for use in other bacteria.

**IMPORTANCE:** *Fusobacterium nucleatum* is an indole-producing human oral anaerobe associated with periodontal diseases, preterm birth, and several cancers. Little is known about the mechanisms of fusobacterial pathogenesis and associated factors mainly due to the lack of robust genetic tools for this organism. Here we showed that a mutated *hicA* gene from *Fuosbacterium periodonticum* expresses an active toxin and was used as a counterselection marker. This *hicA*-based in-frame deletion system efficiently creates in-frame deletion mutations in the wild-type background of *F. nucleatum*. This is the first report to use the *hicA* gene as a counterselection marker in a bacterial genetic study.

## INTRODUCTION

*Fusobacterium nucleatum* is a Gram-negative, indole-producing microorganism naturally isolated from human oral cavity and is especially abundant within the subgingival plaque biofilm composed of more than 400 bacterial species (1, 2). Using three major surface adhesin proteins, FadA (3, 4), RadD (5, 6), and Fap2 (7, 8), *F*. *nucleatum* physically interacts with the host and many other early or late colonizers in oral biofilm, making its presence tightly associated with the onset and development of periodontal disease (2, 9, 10). In addition, evidence has shown that *F. nucleatum* plays an essential role in extra-oral diseases like preterm birth (11, 12) and colorectal cancer (13–16). To better understand the role of this bacterium in various disorders, it is necessary to have comprehensive knowledge of *F. nucleatum’s* biology and pathogenesis. Such knowledge has been dramatically expanded in the past 15 years due to many genetic tools being adapted for use in studies of this organism. There are ways to perform genetic studies of *F. nucleatum*. One way is to use the Tn5-based mutagenesis to produce many random gene mutants in one single assay (7, 17). Another example is to use the defined genetic mutation that makes a single mutant each time. The defined mutations are usually engineered using three methods: 1) plasmid insertion duplication mutagenesis (5, 18), 2) allelic exchange mutagenesis with antibiotic cassette (3), and 3) markerless in-frame deletion (17, 19). The first two methods are easy to carry out and were primarily employed in early studies of *F. nucleatum*. But both have an obvious drawback, frequently creating undesirable polar effects on downstream genes. In addition, the limited number of available selectable markers limits the number of mutations in a strain. However, these issues could be overcome by developing a markerless in-frame deletion system.

The in-frame deletion procedure is a two-step process. First, two similarly sized fragments that are homologous to sequences flanking the target locus are PCR amplified. The two fragments are ligated adjacent to each other in a suicide vector. This construct is subsequently integrated onto the chromosome via homologous recombination and selected using a positive selection pressure, often with antibiotic resistance. Then, the selected simple recombinants are grown without any selective pressure to allow second recombination events and plasmid loss. Next, plasmid-excision/loss recombinants are screened for a second recombination event that occurs between the duplicated homologous chromosomal region which is created after the initial plasmid recombination event. Since the second recombination event is rare, screening these recombinants that have undergone the double crossover is time-consuming. Counterselection strategies are often introduced to facilitate this process to create markerless in-frame deletions (20).

For this purpose, we introduced *galK* as a counterselection marker in *F. nucleatum* (17). The *galK* gene encodes galactokinase, a key enzyme in galactose metabolism. This enzyme phosphorylates D-galactose to generate galactose-1-phosphate and can also convert 2-deoxy-D-galactose (2-DG) into 2-deoxygalctose-1-phosphate, which is not further metabolized and is toxic to the cell. *F. nucleatum* carries a copy of the *galK* gene. To use *galK* for counterselection, a *galK*-minus recipient strain was constructed. An in-frame deletion vector was generated in which the *galK* gene from *F. nucleatum* or *Clostridium acetobutylicum* was cloned, and a strong and constitutive promoter controlled its expression. If the recipient strain lacks galactose kinase (*galK*) and phosphate uridylyltransferase (*galT*), galactose can be used instead of the more expensive 2-deoxy-D-galactose molecule for selection (19). The *galK*-based counterselection system consists of a *galK* minus or *galKT* minus recipient strain and an in-frame deletion vector, which results in a strong negative selection in *F. nucleatum*. This shows that all viable strains when selected with 2-deoxy-D-galactose undergo the second recombination to excise the plasmid. This system successfully generated many fusobacterial in-frame deletions in our lab and other groups (6, 17, 19, 21). However, the apparent limitation of this system is the modified mutant recipient strain requirement. The intermediate product UDP-galactose from galactose utilization is vital in cell envelope maintenance in bacteria (22, 23). The use of galactose present in a culture medium or animal hosts by *F. nucleatum* could change its physiology, as it would potentially module the function of Fap2, an important adhesin that binds galactose and is responsible for cell-cell coaggregation and cell adhesion (7, 8). Consistently, we identified several cases in which the recipient background impacted the phenotypes of our markerless in-frame deletion strains in our studies (unpublished data). Thus, a new counterselection marker is urgently needed to replace *galK* for genetic studies in *F. nucleatum*.

The HicAB is a type II toxin-antitoxin (TA) module that is ubiquitously present in bacteria (24) and is involved in bacterial persistence and chronic infections (25). HicA is a type of ribonuclease that can cause cell growth to stop when expressed without cognate antitoxin HicB. There has only been one study on TA system in the genus *Fusobacterium*, and it is related to a plasmid addition system (Txf-Axf) (26). We tried to explore the possibility of using one of the *hicA* genes from fusobacterial species as a counterselection marker.

In this study, we report the development of a HicA toxin gene-based in-frame deletion system in *F. nucleatum*. We tested this system to construct two markerless in-frame deletions in the fusobacterial adhesin gene *radD* and indole-producing gene *tnaA*. Following counterselection, plasmid loss occurred in every colony tested, while half of the screened isolates grown on plates with theophylline were in-frame deletion mutants. To our knowledge, this is the first report of using a *hicA* as a counterselection in bacteria.

## RESULTS

### A putative mutated HicA toxin from *F. periodonticum* is functional in *F. nucleatum*

In previous studies, we used the *galK* gene as a counterselection marker to efficiently create in-frame deletion mutations in *F. nucleatum* (17). However, the *galK*-based allelic exchange system has an obvious limitation: the host cell must be *galK* minus. Without this gene, bacteria were unable to utilize galactose which is crucial for survival and virulence in certain conditions. This can affect the characteristics of the resulting mutant derivatives. To address this issue, we in this study sought to find a new counterselection marker to replace the *galK* gene for genetic manipulations in *F. nucleatum*.

When beginning our search for a likely candidate, we considered the toxin-antitoxin modules. The toxin-antitoxin systems are widespread in bacteria and typically consist of two components: a toxin that inhibits cell growth and an antitoxin that counteracts the cognate toxin (27). The toxin gene *mazF* in the toxin-antitoxin MazEF modules from *E. coli* has been used for counterselection in various bacteria when *mazF* expression was in highly adjustable controls, often controlled by an inducible promoter (28–33). We first attempted to use a *mazF* as a counterselection marker in *F. nucleatum* but were unsuccessful. Then we switched our attention to the HicA toxin gene in the HicAB toxin-antitoxin module. The HicAB system has been well studied in *E. coli* (34), *Burkholderia pseudomallei* (35) and *Streptococcus mutans* (36). HicA toxins are typically smaller than mazF toxins. MazF in *E. coli* has 111 amino acids and HicA 58. HicA degrades both mRNA and transfer-messenger RNA, halting protein translation and cell growth. MzaF only cleaves mRNA. Co-expression of HicB was able to counteract HicA-induced bacterial growth arrest (36, 37). Since our lab is interested in the biological roles of toxin-antitoxin systems in fusobacterial species, we plan to use an endogenous HicA homolog from the genus *Fusobacterium* and test if it could be employed as a counterselection marker. This way, the codon-bias issue can be efficiently avoided during the ectopic expression of the candidate marker gene in *F. nucleatum*, a low G+C bacterium. Protein expression in a host bacterium is heavily influenced by GC content.

Firstly, we identified *hicA* gene homologs by performing BLAST searches with the *hicA* gene in *B. pesudomallei (BphicA*) as a query against the genome sequence of *Fusobacterium periodonticum*, another oral bacterium. A *hicA* homolog was found in *F. periodonticum* and was named *FpHicA* (see Figure 1A). It encodes a putative 59 amino-acid polypeptide by the gene of *FperA3_010100004605*(https://img.jgi.doe.gov) with 61% identity and 80% similarity to BpHicA. The downstream gene of *FphicA* is the *hicB*, which encodes a putative antitoxin protein for neutralization of HicA toxicity (Figure 1B). Notably, the genome of *F. nucleatum* subsp. *nucleatum* ATCC 23726, our genetic working strain, contains a putative 63 amino-acid peptide HicA homolog (FnHicA) encoded by gene HMPREF0397_0209 (https://img.jgi.doe.gov). However, FnHicA shares only 39% identity and 54%similarity with FpHicA. To use the *FphicA* gene as a counterselection marker in our in-frame-deletion plasmids, the low sequence homology of the *FphicA* gene to the *FnhicA* gene locus would not affect the initial integration of our in-frame deletion constructs.

**Figure 1:**
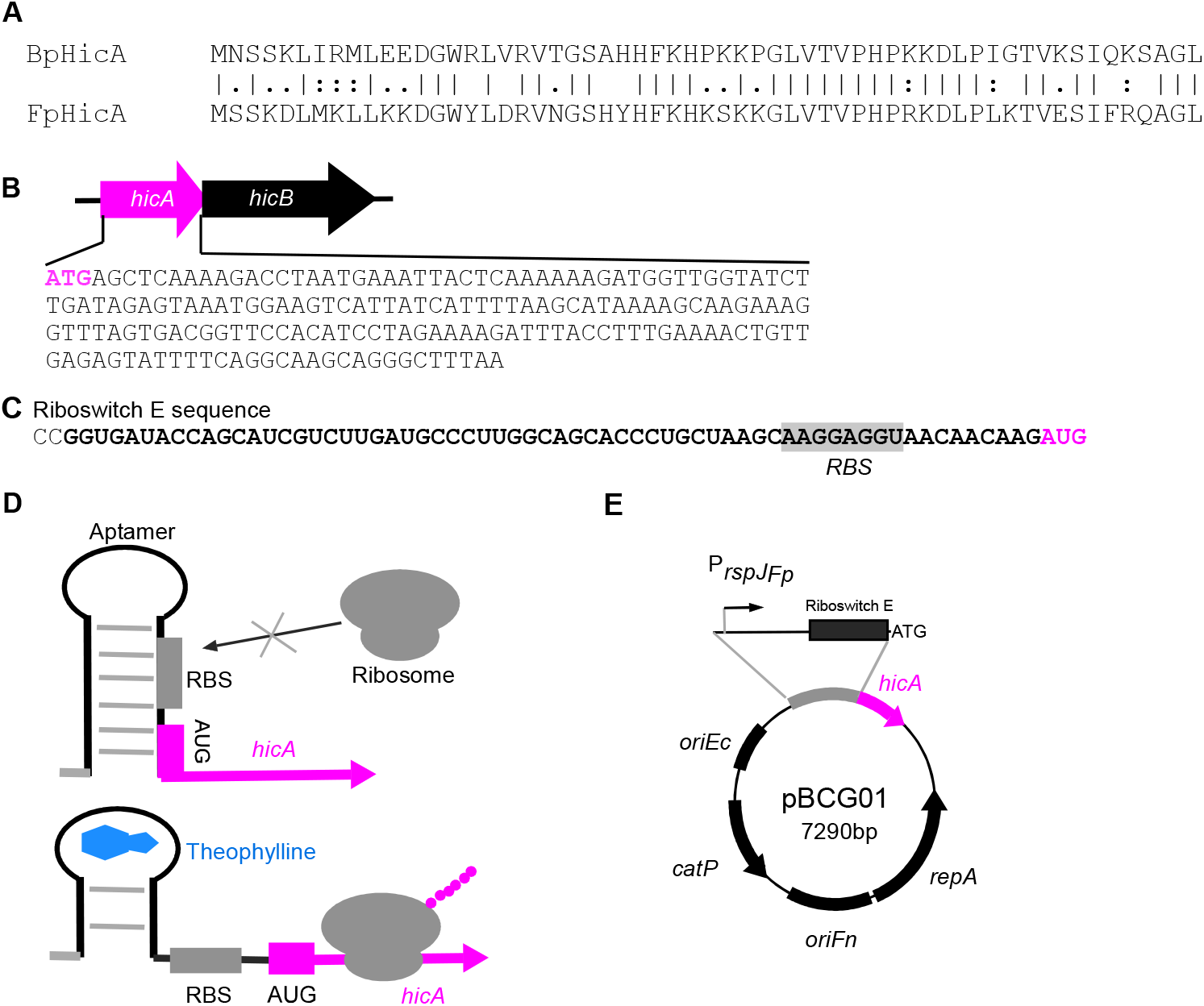
Construction of *E. coli/Fusobacterium* shuttle plasmid pBCG01 expressing putative HicA toxin under the control of riboswitch E-based inducible system. **(A)** Protein alignment of toxin HicA from *Burkholderia pseudomallei* (BpHicA) with HicA encoded by *FperA3_010100004605* (https://img.jgi.doe.gov) in *Fusobacterium periodonticum* ATCC 33693 (FpHicA) using ClustalW program. **(B)** Genetic organization of the *hicA-hicB* toxin-antitoxin pair in *F*. *periodonticum* and the whole DNA sequence of the *hicA* gene. **(C)** The mRNA sequence of the riboswitch E is shown with an aptamer sequence emboldened, ribosome binding site (RBS) sequence highlighted, and start codon red colored. **(D)** Schematic diagrams of a functional model of the theophylline responsive riboswitch. Without theophylline, the riboswitch forms a stem-loop structure that conceals the RBS in the mRNA transcript. The riboswitch alters its confirmation when theophylline binds to the aptamer, resulting in the release of the RBS and the initiation of translation of *hicA*. **(E)** *E. coli/Fusobacterium* shuttle plasmid pBCG01 containing *hicA* gene under the control of *rpsJ* promoter from *F*. *periodonticum*. The RBS in the *rpsJ* promoter region was replaced by the riboswitch E element. ColE1 RNA II origin (*ori_Ec_*), the pFN1 ori (*ori_Fn_*), *repA* gene, and *catP* gene are indicated. The *ori_Ec_* allows replication of the shuttle plasmid in *E. coli*, and the plasmid replicates in *F. nucleatum*, requiring the *ori_Fn_* and *repA*.

Next, we wanted to express the *hicA* gene in *F. nucleatum* ectopically and test the toxicity of the FpHicA protein. To do so, the Gibson assembly cloning was performed to construct a pCWU6-based (17) plasmid that expresses the *hicA* gene. The *rpsJ* gene promoter was proven as a strong, constitutive promoter in *F. nucleatum* (6). Based on this finding, we hypothesized that the promoter of the *rpsJ* gene in *F. periodonticum* (FperA3_010100002588, https://img.jgi.doe.gov/) would be similar. As a result, we used the *rpsJ_Fp_* promoter to drive *hicA* expression. To control the putative HicA expression level, a well-characterized synthetic theophylline-responsive riboswitch E (38) (Figure 1C) was inserted in between the *rpsJ_fp_* promoter and the *hicA* start codon (Figure 1E). This riboswitch E unit is functional in Gram-positive and Gram-negative bacteria (38, 39). It is comprised of an aptamer and a synthetic ribosome binding site (RBS). Theoretically, transcription of the riboswitch under the control of the *rpsJ_Fp_* promoter occurs constitutively during cell growth. However, the synthetic RBS located downstream of the aptamer is sequestered via pairing with the stem of this riboswitch, resulting in a translational stop. The binding of ligand theophylline exposes the RBS to the translation machine by altering the downstream base pairing. Consequently, the *hicA* gene translation begins when the ribosome binds to the RBS (Figure 1D). We wanted to obtain a plasmid that ectopically expresses the *hicA* gene under the control of the riboswitch E element. Four positive *E. coli* clones generated by the Gibson assembly procedure were randomly chosen to culture for plasmid isolation. Before the four plasmids were verified by DNA sequencing, we transformed them individually into *F. nucleatum* ATCC 23726 strain by electroporation. Cells were then plated on agar plate with thiamphenicol. The colonies formed after three days of growth in an anaerobic chamber. One colony from each plasmid transformation plate was picked up to grow in the TSPC medium. The growth of these transformants was similar to the wild-type strain when *hicA* gene expression was not induced with theophylline (data not shown). Interestingly, only one transformant did not grow when 2 mM theophylline was provided—the other three grew the same with or without the theophylline (Figure 2C and data not shown). Subsequently, DNA sequencing revealed that the three plasmids are identical and each contains a complete *hicA* gene. Surprisingly, the plasmid conferring cell growth arrest had a point mutation in a “4A” sequence (Figure 2A). There were no errors in the promoter region, riboswitch E unit, or other parts of the *hicA* gene in the four sequenced plasmids. An adenine base missed in the “4A” sequence (Figure 2B), subsequently refered to as *hicA(3A*), which caused a frameshift and introduced a premature stop codon at codon 6. It presumably produced a five-amino acid peptide (MSSKT) starting its translation from the first ATG codon when the inducer was added. By closely examining the mutated *hicA* sequence, we found that the introduced stop codon TAA had one nucleotide overlapping with a second ATG codon: codon 7 of the wild-type *hicA* gene (Figure 2A). This ATG at codon seven is postulated to allow *hicA(3A*) mRNA to restart its translation, creating a shorter HicA protein lacking the first six amino acids (Figure 2B). In other words, it is possible that the production of truncated HicA protein is functional in *F. nucleatum* and able to cease cell growth, but not its full-length version. As the produced MSSKT peptide is a small peptide, it may act as anantimicrobial peptide (AMP) that kills or prevents bacterial growth (40). To exclude the possibility of the MSSKT peptide acting as an AMP, the gene region only synthesized with the five-amino peptide was amplified from the mutated plasmid pBCG02 and re-cloned in pCWU6. Its expression was under the control *rpsJ_FP_* promoter and the riboswitch E unit (pBCG04), the same as those in pBCG01 and pBCG02. Expectedly, in the presence of inducers, the expression of the short peptide did not affect bacterial growth (Figure 2C, pBCG04). The *hicB* gene could counteract the toxicity of mutated *hicA(3A*). As the *hicB* was co-expressed with mutated *hicA*, fusobacterial growth arrest was reversed (Figure 2C, pBCG03) to normal levels when grown on agar plates with 2 mM theophylline (Figure 2C). We were interested in knowing if the mutated *hicA* gene was functional in its native host, *F*. *periodonticum* as well. For that purpose, the pBCG02 was transformed into *F*. *periodonticum* by electroporation, and one transformant was further cultured and streaked on the plates. In the presence of 2 mM inducer theophylline, cell growth was stopped entirely. In contrast, the cells of *F*. *periodonticum* with pBCG01 containing the wild-type *hicA* gene showed no signs of a growth defect when *hicA* gene expression was induced (Figure 2D). In addition, we showed that induction of the mutated *hicA* gene prevented cell growth of *E. coli* in the presence of 2 mM theophylline (Figure 2E). Together, these data suggested that the putative mutated HicA with 53 amino acids functions as a toxin in our tested bacteria.

**Figure 2:**
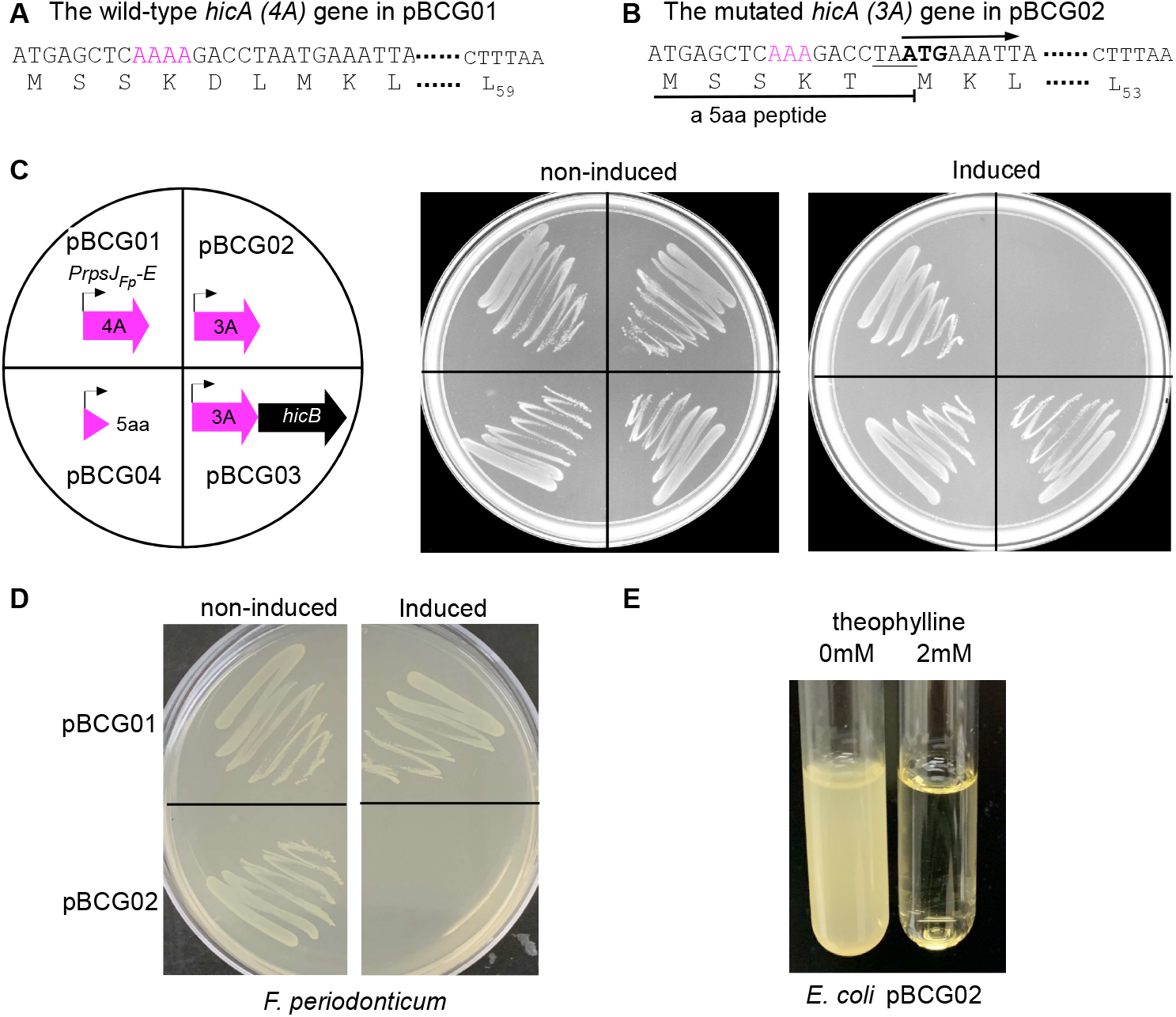
A putative shorter HicA protein that lacks the first six amino acids is toxic in *Fusobacterium nucleatum* and other bacteria. **(A)** The plasmid pBCG01 contains an intact *hicA* gene, indicating the presence of a four consecutive adenine base sequence (4A) after the start codon (red colored). The wild-type *hicA* putatively encodes a 59-amino acid protein. **(B)** An adenine base was missing from the 4A sequence resulting in the *hicA* gene (3A) during construction of pBCG01. We later named that mutated construct as pBCG02. This point mutation causes a frameshift resulting in a premature stop codon (TAA, underlined) at codon six and generates five amino acids peptide (MSSKT). The *hicA(3A)* mRNA then restarts its translation from the new initiation site (ATG, emboldened), giving a putative shorter HicA protein lacking the first six amino acids. **(C)** Growth of *F. nucleatum* strain 23726 harboring the plasmid construct pGCG01, pBCG02, pBCG03, or pBCG04 on TSPC agar plates supplemented with thiamphenicol plus 0 (non-induced) or 2mM (induced) theophylline and incubated at 37°C in an anaerobic chamber for two days. Schematic representations of the constructs are shown in the left diagram. **(D)** Growth of *F. periodonticum* containing the plasmid pBCG01 or pBCG02 on TSPC agar plates under the uninduced or induced condition with theophylline after two days of growth in an anaerobic chamber. **(E)** Overnight E. *coli* containing pBCG02 was diluted by 1:1000 and aliquoted into two tubes with 6 ml LB broth plus an indicated concertation of theophylline after 18 hours of growth with shaking (200 rpm) at 37°C. These experiments were performed more than three times, and a representative result was presented.

### Use of mutated *hicA* as a counterselection marker for in-frame deletion in *F. nucleatum*

The ectopic expression of the putative HicA(3A) inhibited fusobacterial cell growth, indicating that *hicA (3A*) gene might be suitable for counter-selection in *F. nucleatum*. To test this possibility, an allelic exchange vector named pBCG02-Δ*radD* (Figure 3A) was constructed to delete the gene region coding for the amino acids in positions between 44 and 1240 of the RadD protein. Our data indicated that the RadD (44-1240) region contains a motif critical for RadD activity (unpublished data). The RadD protein is a large surface adhesin that mediates *F. nucleatum’s* physical interactions with some oral bacteria, mostly Gram-positive, early colonizers in the human mouth (5). These aggregated interactions are critical for dental plaque formation. An inverse PCR linearized the pBCG02 with primer pair pCWU6-F/R to remove *repA* and *ori_FN_* (Figure 3A). The RepA and *ori_FN_* elements are required for pCWU6-based plasmid replication in *F. nucleatum* (18). Without them, the shuttle plasmid pBCG02 becomes a suicide vector with a positive selection marker (*catP*) and a negative selection marker (*hicA(3A)*). The same length of upstream and downstream DNA sequences flanking the region -responsible for the synthesis of the RadD (44-1240)- was amplified. An overlapping PCR reaction then fused the upstream and downstream fragments. Gibson assembly was used to clone the fused fragment into the linearized pBCG02 that lacks *repA* and *ori_FN_*. The resulting in-frame deletion construct was transformed into *E. coli* C2987. Positive colonies were confirmed by colony PCR using the primer pair Δ*radD-*F/R. Positive colonies showed a 1.34 kb insert and were further selected for plasmid preparation and sequencing. The transfer of the allelic exchange plasmid pBCG02-Δ*radD* into *F. nucleatum* ATCC 23726 was traditionally performed via the electroporation procedure that our lab used previously (41). The selection of recombinant vectors to integrate into the bacterial chromosome was performed by thiamphenicol, a chloramphenicol derivative. One randomly selected thiamphenicol-resistant integration colony was chosen for theophylline-negative selection based on putative HicA(3A) toxicity. This strain was grown in the absence of antibiotics overnight and then diluted by 1 to 1000 the following day before plating cells on the TSPC agar plates with 2 mM theophylline. To estimate the frequency of plasmid excision, a plate without an inducer was used to grow the un-counter-selected cells as a control. As shown in Figure 3B, the theophylline-negative selection only allowed a small fraction of the cells to survive. Based on the ratio of theophylline-resistant cells to total cells (cells grown on plates without selection, see Figure 3B, the left plate without an inducer), we estimated that approximately 1 in 2000 cells underwent a second recombination to lose the integrated plasmid. Eight randomly selected theophylline-resistant colonies were confirmed for antibiotic sensitivity to screen for possible in-frame deletion mutants. Next, colony PCR was used to detect their genotypes using the primer pair Δ*radD-*F/R illustrated the Figure 3A. As shown in Figure 3C, Four of the eight colonies generated amplicons of approximately 1.4 kb, indicating an in-frame deletion. The remaining four colonies produced the expected wild-type amplicons of approximately 5.0 kb. The coaggregation assay was also used to confirm the four generated *radD* mutant strains and four wild-type revertant strains. We expected coaggregation defects in these mutant strains because this region of RadD (44-1240) is located in the passenger domain, which is responsible for *F. nucleatum* coaggregation with oral bacterium *Actinomyces oris* (6). This was indeed the case (Figure 3D). Clones 1, 2, 4, and 8 showed coaggregation defects with *A. oris*, whereas all wild-type reverants (clones 3, 5, 6 and 7) exhibited aggregation similar to the wild-type strain. Finally, two of the in-frame deletion mutant clones (clone 1 and 4) were also sequenced and found to exhibit the expected in-frame deletion that the *radD* gene only lacks in the desired deletion region by amplicon sequencing.

**Figure 3:**
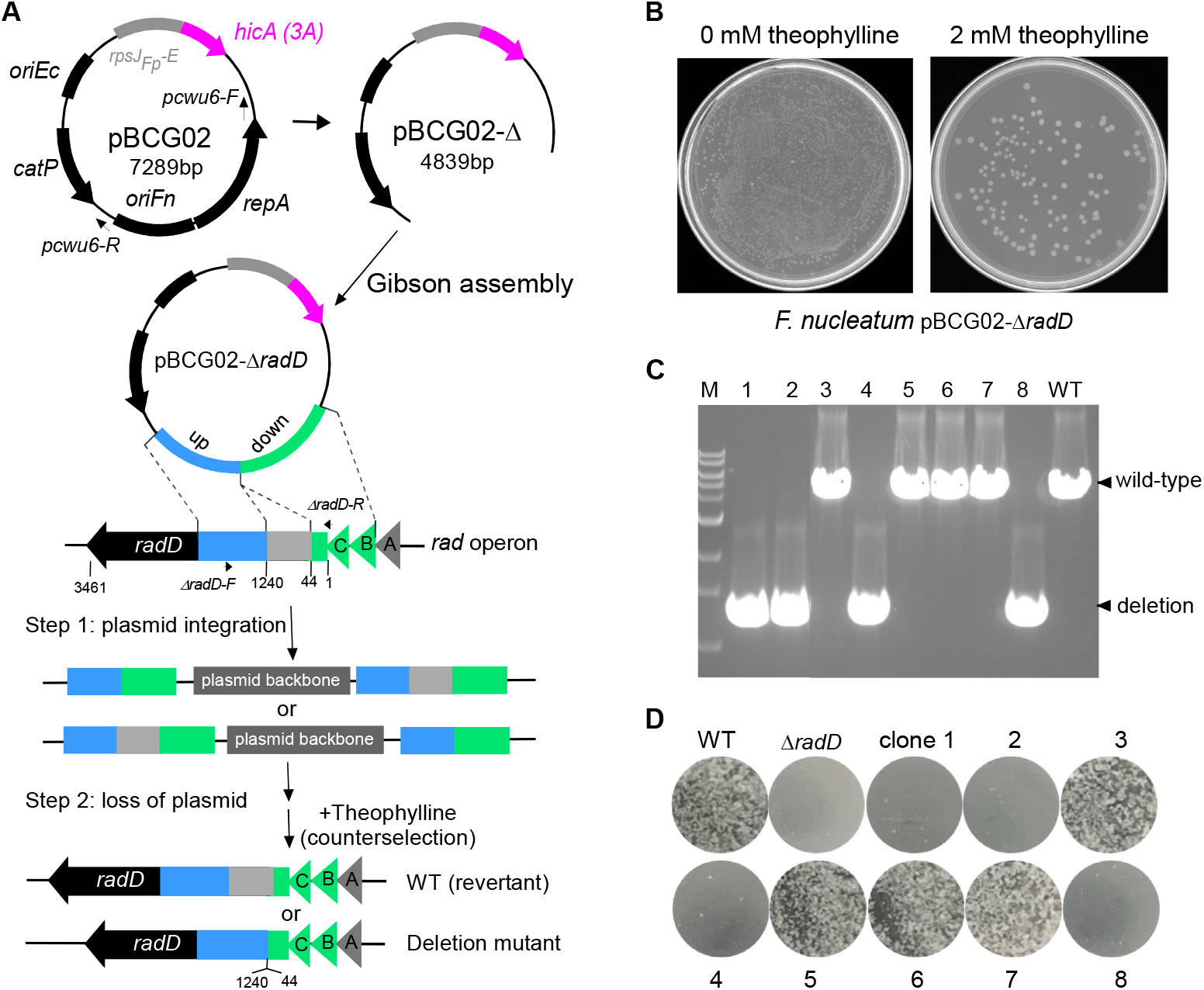
Applying *hicA(3A)* as a counter-selectable marker for gene deletion in *F. nucleatum*. **(A)** Graphic representation of producing the non-replicative vector backbone pBCG02-Δ used to generate unmarked, in-frame gene deletions in *F. nucleatum* via a two-step strategy. The primer pairs (pCWU6-F/R, indicated by short arrows) were used to perform reverse PCR to remove the region, including the *ori_Fn_* and *repA*. The *radABCD* operon is shown. Approximately 1.6 kb fragments homologous to the upstream (blue) and downstream (green) regions of the designed deletion region in *radD* were amplified by PCR. They have been ligated adjacently on the vector backbone pBCG02-Δ via the Gibson assembly procedure. The resulting plasmid pBCG02-Δ*radD* was used to generate an unmarked *radD* deletion (the gene deletion region coding for the amino acids in positions between 44 and 1240 of the RadD protein of 3461 amino acids) mutant in the wild-type background by a two-step allelic exchange. **(B)** TSPC agar plate representative of the counterselection effect by putative HicA(3A) toxin expression in *F. nucleatum* A 100 μl aliquot of 1,000-fold diluted overnight culture for pBCG02-pBCG02-Δ*radD* integrated strain was plated on TSPC agar plates containing 0- or 2-mM inducer theophylline to select for cells in which the plasmid had been excised and lost. **(C)** The screen of *radD* deletion mutants by colony PCR with the primer pair Δ*radD*-F/R (indicated in **A**). Lane 1-8, 10 μl of PCR products of eight colonies randomly selected from the TSPC agar plate. Four of the eight chosen random colonies were deletion mutants, while four reverted to wild-type during the second recombination event. Bands of approximately 1.4 kb indicate *radD* deletion, whereas bands of about 5.0 kb indicate a wild-type (WT) genotype. **(D)** *F. nucleatum* wild-type, *radD* mutant strain and 8 counterselection isolates from (C) were examined for coaggregation with *A. oris* MG-1. A representative result was presented after these experiments were carried out three times.

To further test this *hicA(3A)-based* counterselection system, we used it to make an in-frame deletion of the *tnaA* gene (HMPREF0397_1802, https://img.jgi.doe.gov/) in *F. nucleatum*. It has been reported that the *tnaA* gene encodes the enzyme tryptophanase that produces indole from tryptophan in *E. coli* (42). As a signaling molecule, indole plays an essential role in bacterial physiology, ecological balance, and human health (43). However, its biological function in *F. nucleatum* has yet to be explored. An allelic exchange cassette carrying the flanking regions of *tnaA* was generated by an overlapping PCR. The allelic exchange cassette was cloned into the linearized pBCG02, in which *repA* and *ori_Fn_* were omitted. The generated suicide plasmid pBCG02*-*Δ*tnaA* (Figure 4A) was then introduced into an *F. nucleatum* wild-type background strain by electroporation. The insertion obtained by homologous recombination of the plasmid into the chromosome was selected in the presence of thiamphenicol. A thiamphenicol-resistant colony was grown in TSPC without antibiotics overnight. The following day, aliquots of 1000-fold diluted cultures were plated on TSPC agar plates containing 2 mM theophylline. After three days, ten theophylline-resistant colonies were randomly chosen to check thiamphenicol sensitivity (100% efficiency). The colony PCR was then used to analyze the genotypes of the ten colonies with primer pair TnaAupF and TnaAdnR (see Table 2) indicated in Figure 4A. The results revealed that half of the colonies were the expected deletion mutant strains, and half were the wild-type revertant strain (Figure 4B). Lastly, the *tnaA* deletion mutant was confirmed using the indole assay, which revealed that the wild-type cell and wild-type revertant cell (clones 4, 6,7, 8, and 10) developed red color, whereas the *tnaA* mutant (clones 1, 2, 3, 5, and 9) did not (Figure 4C). This suggests that the *tnaA* homolog in *F. nucleatum* is responsible for indole production.

**Figure 4:**
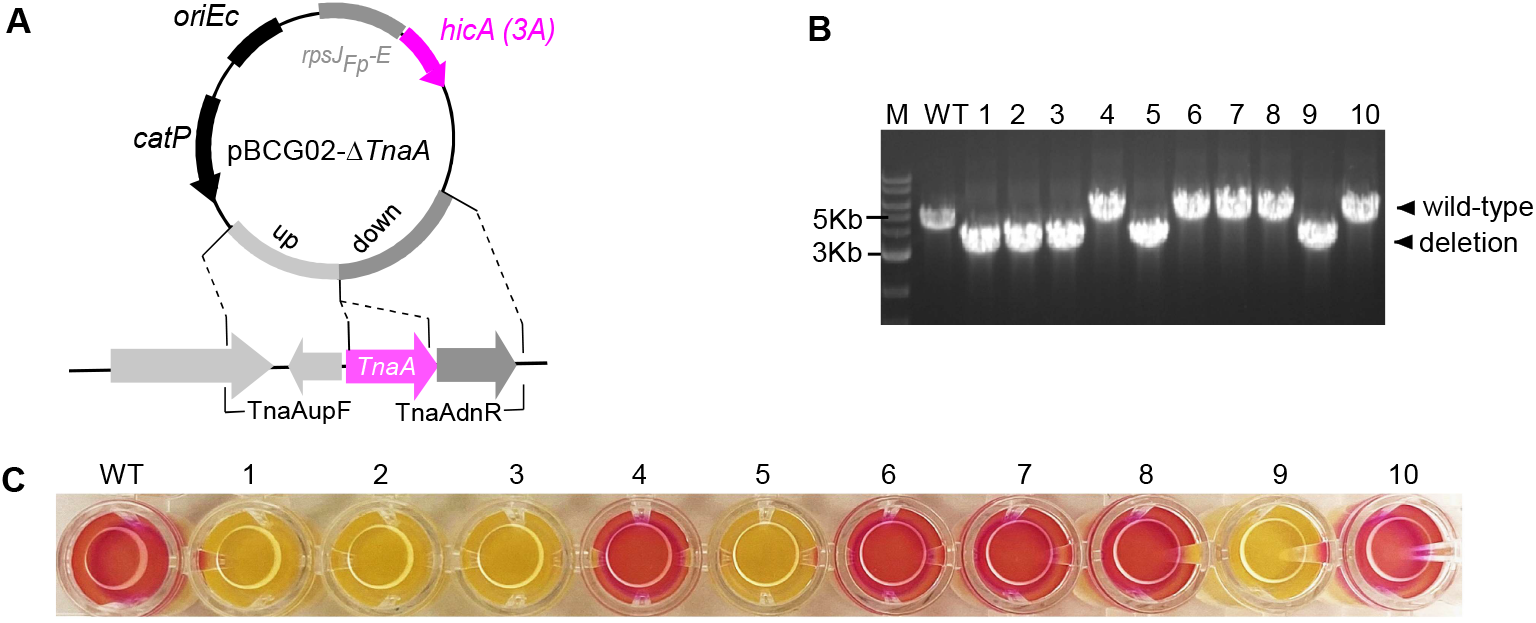
Deletion of indole-producing gene *tnaA* in *F. nucleatum* as *hicA(3A)* as a counter-selectable marker. **(A)** Schematic drawing of pBCG02-Δ*tnaA* for the deletion of tryptophanase TnaA gene. Approximately 1.6-kb fragments homologous to the upstream (blue) and downstream (green) regions of the *tnaA* gene were ligated adjacently on the vector backbone pBCG02-Δ via Gibson assembly procedure to create the vector pBCG02-Δ*tnaA*. The primer pair-for colony PCR to screen Δ*tnaA* mutant strains were indicated by small arrows. **(B)** The genotype of *tnaA* mutant strains following *hicA(3A*) counterselection. Bands of approximately 3.2 kb indicate a *tnaA* deletion, whereas bands of about 5.0 kb indicate a wild-type genotype. “WT” represents the *F. nucleatum* wild-type. **(C)** *F. nucleatum* wild-type and 10 isolated colonies after counterselection from (B) were examined for indole production. Indole production was demonstrated by adding Kovac’s reagent (100 μl) into a 200 μl of individual culture in a 96-well plate, which reacts with the indole giving a red color in wild-type genotype fusobacterial cultures. Three repeats of the studies were conducted, and one typical outcome is shown.

**Table 1:** Bacterial strains and plasmids used

| Strain & Plasmid | Description | Reference |
| --- | --- | --- |
| Strains |  |  |
| *F. periodonticum* 33693 | Parental strain | This study |
| *F. nucleatum* 23726 | Parental strain | (5) |
| *F. nucleatum* BG01 | $\Delta radD$; an isogenic derivative of 23726 | This study |
| *F. nucleatum* ZP01 | $\Delta tnaA$; an isogenic derivative of 23726 | This study |
| *A. oris* MG1 | Wild Type strain | (48) |
| *E. coli* C2987 | Cloning host | From NEB |
| Plasmids |  |  |
| pCWU6 | *E. coli*/*Fusobacterium* shuttle vector, chloramphenicol /thiamphenicol resistance; $cm^R$/thia<sup>R</sup> | (18) |
| pRECas2 | The J23119 promoter incorporating the riboswitch element E controls Cas9 expression | (39) |
| pBCG01 | pCWU6 expressing wild-type *hicA* gene [*hicA*(4A)] from *F. periodonticum* 33693 under the control of the *rpsJ<sub>FP</sub>* promoter and its ribosome binding site was replaced by a riboswitch element E from pRECas2 | This study |
| pBCG02 | Derivative of pBCG01 expressing *hicA* with a point mutation [*hicA*(3A)]. The *hicA*(3A) mRNA is translated from the initiation site met-7, giving a shorter protein lacking the first six amino acids. | This study |
| pBCG03 | Derivative of pBCG02 co-expressing *hicA*(3A) and *hicB*, the cognate *hicA* anti-toxin gene | This study |
| pBCG04 | Derivative of pBCG02 expressing the five amino acids peptide (MSSKT) alone | This study |
| pBCG02-$\Delta radD$ | Derivative of pBCG02 lacking *repA* and *ori<sub>Fn</sub>*, deletion vector of *radD* $\Delta 41-1240$ | This study |
| pBCG02-$\Delta tnaA$ | Derivative of pBCG02 lacking *repA* and *ori<sub>Fn</sub>*, deletion vector of *tnaA* | This study |

**Table 2:**
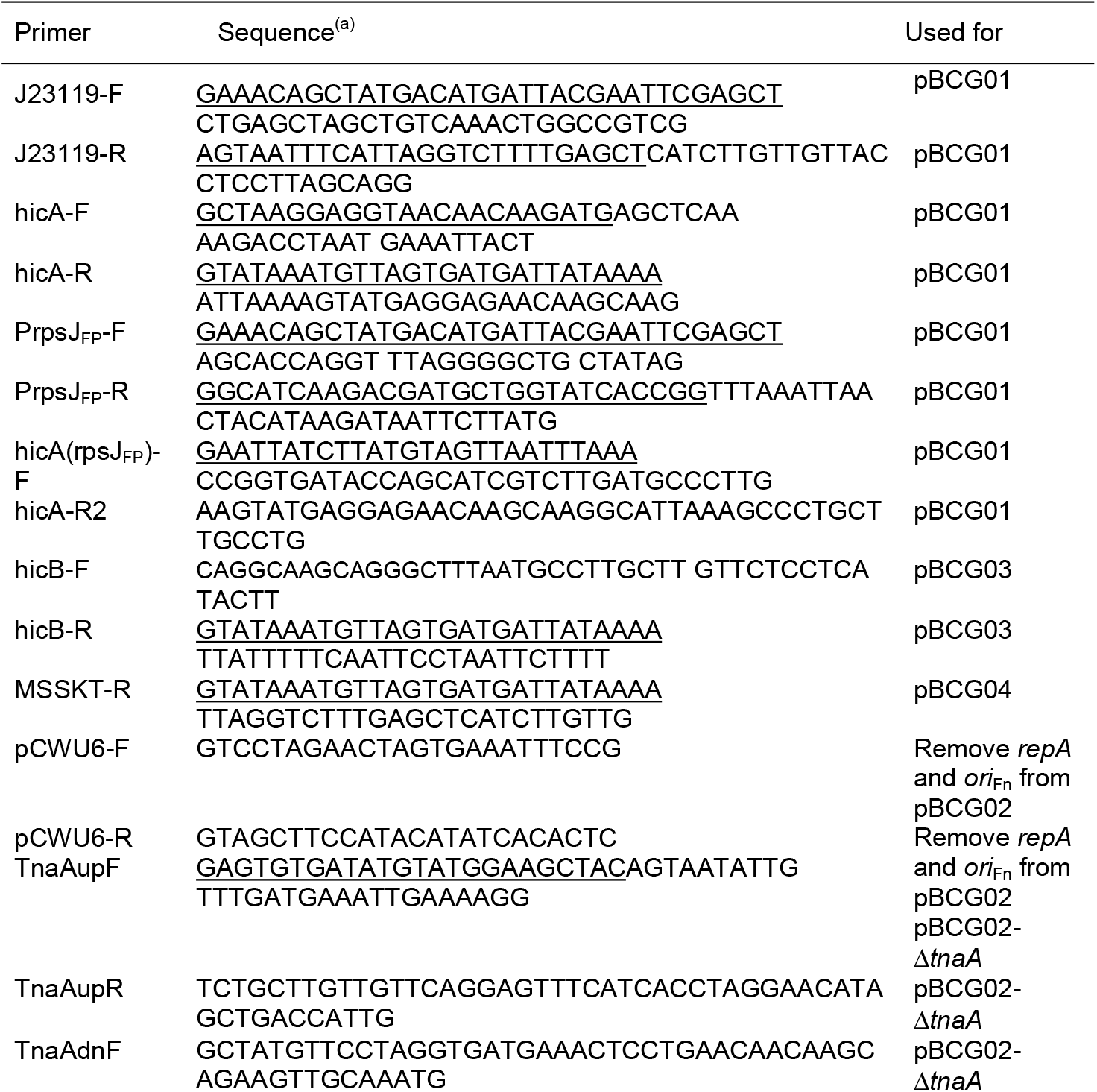
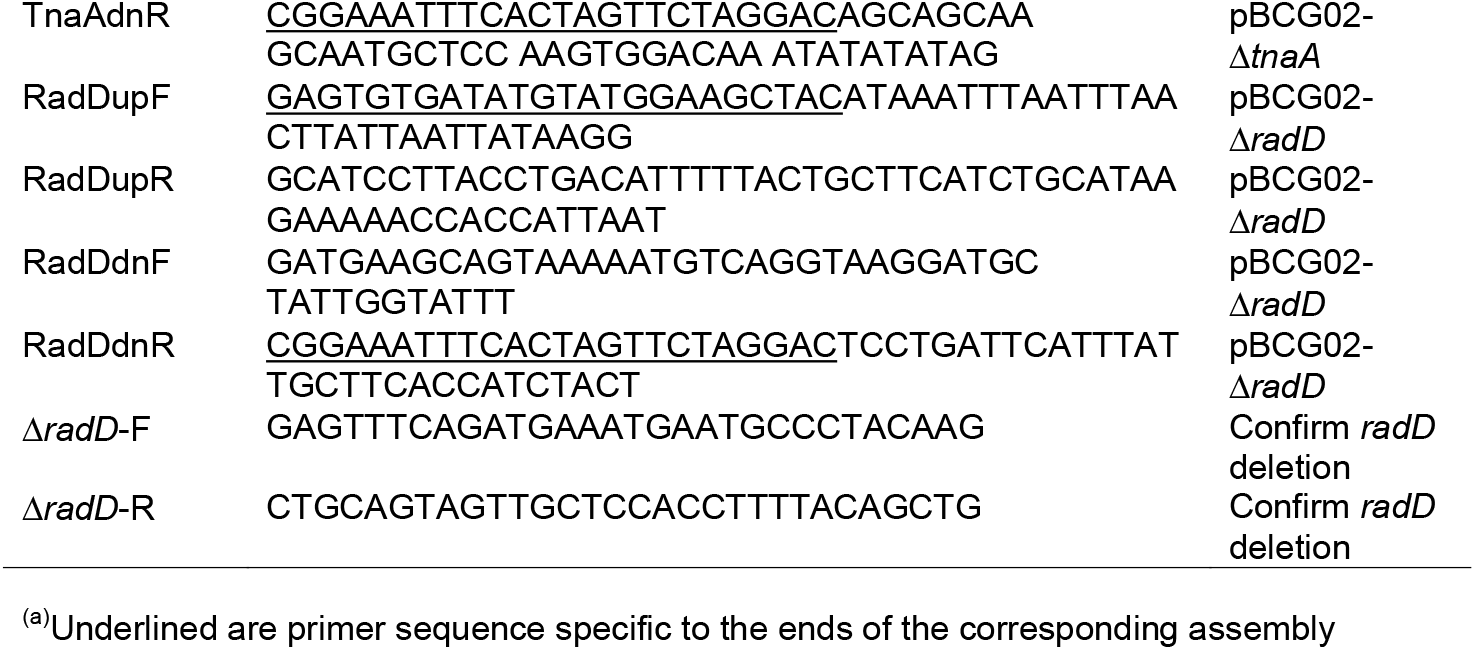
Primers used in this study

## DISCUSSION

The HicAB toxin-antitoxin system has been widely studied in bacteria. However, no HicA toxin has been used for counterselection to create markerless in-frame deletion mutations. HicA is a ribonuclease, and its expression prevents cell growth in many bacteria (24, 34). In this study, we used a *hicA* gene from *F*. *periodonticum* and established a facile in-frame deletion system in *F. nucleatum*. The success of establishing this putative HicA toxin for counterselection in *F. nucleatum* has benefited from one unexpected finding. The expression of a full-length *hicA* gene, encoded for a putatvie 59 amino-acid HicA toxin, did not induce cell death; however, when inducing expression of a short form of putative HicA protein that lacks the first six amino acids, the cells died (Figure 2C). In other words, the putative truncated version of HicA possesses a toxin activity. The finding of this active form of putative HicA was accidental. An unexpected point mutation in *hicA* occurred during plasmid constructions, allowing *hicA* mRNA translation to start from the second ATG, codon 7, in its original version of this gene (Figure 2B). The presence of N-terminal six amino acids in the putative HicA inhibits its activity. Some proteins, like bacteriocins, must remove part of their N-terminus before being active (44, 45). For toxin proteins from all studied toxin-antitoxin systems, there is no such report that the removal of partial N-terminal sequences is needed for their activities. *F*. *periodonticum* may produce a nonfunctional premature HicA protein, which then requires an unknown mechanism to cleave its N-terminal partial sequence in order to be in an active state. Consistently with this possibility, the *hicA* expression in the *hicB* deletion mutants does not exhibit growth defects in *E. coli* (46), *Haemophilus ducreyi* (47), and *Streptococcus mutans* (36). Another possibility is that the translation of this gene in *F*. *periodonticum* may start from the second ATG codon -the codon 7 of the gene annotated in the genome database (https://img.jgi.doe.gov)-but not the first ATG as its starting codon. All of these possibilities require further studies.

This work aimed to find a new gene to replace *galK* as a counterselection marker to generate markerless gene deletions in *F. nucleatum*. We showed that cell growth would be arrested entirely when the mutated *hicA* was induced in the wild-type background of *F. nucleatum* (Figure 2C). So, we attempted to use the truncated form of *hicA* gene as a counterselection maker in allelic exchange constructs for creating in-frame-deletion mutations in *F. nucleatum*. The *rpsJ* promoter is a constitutive and robust promoter in many bacteria (6, 48). We chose to use the *rpsJ* promoter from *F*. *periodonticum* to drive the expression of the putatively active HicA toxin. Choosing the promoter from *F*. *periodonticum*, but not *F. nucleatum*, was to avoid the potential interference to the initial integration of our constructs due to the homology of this promoter fragment to the native *rpsJ* locus. A well-studied theophylline-responsive synthetic riboswitch was employed to control the active *hicA* gene expression. Six synthetic theophylline-responsible riboswitches were widely used in various bacteria (38, 39, 49). The riboswitch E was chosen to control *hicA* mRNA translation because only this riboswitch works and does not have a leaking issue when used in *F. nucleatum* (unpublished data). This theophylline-sensing riboswitch tightly controls the expression of the active HicA protein. Since 2 mM of theophylline could induce enough of the toxic protein to prevent cell growth, no step was carried out to optimize theophylline’s concentration further. With this system, we successfully created the in-frame deletion for the fusobacterial adhesin protein RadD gene and the *tnaA* gene responsible for indole production in *F. nucleatum*.

The *hicA(3A)*-based system showed a reliable and robust counterselection exactly like *galK*-utilized counterselection. All colonies were thiamphenicol-sensitive after counterselection, which indicated that 100% of colonies underwent loss of the integration plasmids once the putatively active HicA toxin protein was induced by 2 mM theophylline. Compared with the *galK* system, the *hicA(3A)*-dependent counterselection has an immediate advantage: it can work in the wild-type background of *F. nucleatum*. MazF has been used as a counterselection marker in other Gram-positive and Gram-negative bacteria (20). As we prepared this manuscript, a MazF-mediated counterselection strategy was reported in *F. nucleatum* (50). They use a tetracycline-inducible system to control the MazF expression in *F. nucleatum*. The expression of MazF is lethal. That allowed the counterselection in *F. nucleatum* due to MazF toxicity, and only cells lacking the integrated plasmid could grow. Undoubtedly, this system is excellent and is easy to use in fusobacterial genetic study. But, it has one potential problem: many strains of *F. nucleatum* are tetracycline sensitive. We have found that even 20ng/ml tetracycline or 80ng/ml anhydrotetracycline discourages the growth of *F. nucleatum* ATCC 23726, and some are even more sensitive. That may limit the use of the tetracycline-induced MazF system for counterselection in some strains of *F. nucleatum*. In this aspect, the theophylline-sensed riboswitch is better than the tetracycline-inducible system since *F. nucleatum’s* growth is not affected by the presence of theophylline, even at high concertation (data not shown).

The results presented in this study support the use of HicA toxin activity for negative selection in *F. nucleatum*. The active HicA may work in many bacteria, as we already demonstrated that the induction of this putatively active HicA causes cell death in *F*. *periodonticum* and *E. coli* (Figure 2D, 2E). As far as we know, this is the first time the *hicA* gene as a counterselection marker for bacterial genetic studies. This same strategy is also theoretically feasible in other bacteria that do not already possess an efficient negative selection system. Currently, we are working on this system in *F. periodonticum* to test if it can be widely applied in other fusobacterial species.

## MATERIALS AND METHODS

### Bacterial strains, plasmids, and growth conditions

Bacterial strains and plasmids used in this study are listed in Table 1. *F. nucleatum* and *F. periodonticum* strains were grown in tryptic soy broth (TSB) supplemented with 1% Bacto^™^ peptone plus 0.25% freshly made cysteine (TSPC) or on TSPC agar plates in an anaerobic chamber filled with 80% N_2_, 10% H_2_ and 10% CO_2_. Hemin (5 μg/ml) and vitamin K3 (1 μg/ml) were added for *F. periodonticum* growth on TSPC agar plates. *A. oris* were grown in Heart Infusion Broth (HIB). *E. coli* strains were grown in Luria-Bertani (LB) broth. Fusobacterial transformants with pBCG01, pBCG02, pBCG03, or pBCG04 were grown overnight in TSPC medium with 5 μg/ml thiamphenicol. Overnight cultures were directly streaked on TSPC agar with antibiotic selection, with or without 2 mM theophylline, after 72 h of incubation at 37°C in an anaerobic chamber. 40 mM theophylline stock was prepared in a TSP medium. pREcas2 was purchased from Synthetic Biology Research Center at University of Nottingham, UK. Antibiotics used as needed were: chloramphenicol (15 μg ml^-1^) and thiamphenicol (5 μg ml^-1^). Reagents were purchased from Sigma unless indicated otherwise.

### Plasmid construction

All plasmids used in this study were constructed via Gibson assembly cloning, essentially performed according to the manufacturer’s instructions with NEBuilder^®^ HIFI DNA Assembly Master Mix. This method requires adjacent segments with at least 18-bp long, identical sequences on the ends. These identical sequences were generated via PCR with primers that contain a 5’ end identical to an adjacent segment and a 3’ end that anneals to the gene-of-interest sequence using a 2x hot-start PCR master mix (Takara, #R405A). The primers for PCR reactions were listed in Table 2.

pBCG01 and pBCG02– To create these plasmids, pREcas2 (39) was used as a DNA template to amplify the J23119 promoter region and the riboswitch E element with primers J23119-F/R. The primer pair hicA-F/R (Table 2) was used to PCR-amplify out the *hicA* gene (see gene identification no. FperA3_010100004605 in the *F. periodonticum* database at https://img.jgi.doe.gov/) with genome DNA of *F. periodonticum* ATCC 33693 as a template. The two amplicons were fused via an overlapping PCR reaction, and the resulting PCR product was designated as *J23119-E-hicA*. Then the fragment encompassing the riboswitch E and *hicA* (E-hicA) was amplified with primers hicA(rpsJ_FP_)-F and hicA-R. The primer pair PrpsJ_FP_-F/R was used to amplify the promoter region of the *rpsJ* gene (FperA3_010100002588, https://img.jgi.doe.gov/) without its ribosome binding site (PrpsJ_FP_) with the genome DNA of *F. periodonticum* ATCC 33693. The *E-hicA* and *PrpsJ_FP_* amplicons were cloned into SacI and HindIII-digested pCWU6 (17) by Gibson assembly and transformed into *E. coli* C2987 cells, as described previously (51). Four positive clones were chosen to culture for plasmid isolation, and the four plasmids were sent for sequencing. Three had the wild-type *hicA* DNA sequence, but one had a point mutation at the “4A” sequence (see Figure 2A, 2B). The plasmid that contains this point mutation was named pBCG02.

pBCG03 –To test if the putative antitoxin HicB prevents the putative HicA (3A) toxicity in *F. nucleatum*, the *hicB* gene (FperA3_010100004610, https://img.jgi.doe.gov/) was placed at immediate downstream of *hicA*, and the expression of the two genes was under the control of *rpsJ_FP_* promoter and riboswitch E unit. The fragment, including *PrpsJ_FP_-E-hicA*, was amplified with the primers PrpsJ_FP_-F and hicA-R2 from pBCG02. Secondly, the primer pairs hicB-F/R were used to PCR amplify the *hicB* gene. The two amplicons of *PrpsJ_FP_-E-hicA* and *hicB* were mixed with SacI/HindIII-cut pCWU6 in the Gibson assembly master mix solution to generate pBCG03.

pBCG04 – With plasmid pBCG02 as a DNA template, the primer pairs PrpsJ_FP_-F and MSSKT-R were used to amplify the fragment encompassing PrpsJ_FP_-E and *hicA* (3A) region only, including the first ATG to the premature stop codon due to missing one “A” base in “4A” sequence. The short *hicA* region codes a five amino-acid peptide (MSSKT, see Figure 2B). This amplicon was cloned into pCWU6 precut by SacI/HindIII via Gibson assembly to generate pBCG04.

All produced plasmids were confirmed by sequencing and then transformed into *F. nucleatum* ATCC 23726 or *F*. *periodonticum* ATCC 33693 with the electroporation procedure we previously published (41). In brief, cells of either *F. nucleatum* or *F*. *periodonticum* were harvested by centrifugation from a 100 ml stationary phase culture. The cells were washed twice with sterile water and once with 10% glycerol, then resuspended in 3 ml of 10% glycerol and divided into 0.2 ml aliquots. Next, 1.0 mg purified plasmid DNA was added to 0.2 ml aliquot of the above-prepared fusobacterial cells in a pre-cooled corvette (0.1 cm electrode gap, Bio-Rad) and left on ice for 10 minutes. The reactions were electroporated using a Bio-Rad Gene Pulser II set at 25 kV/cm, 25 F, and 200 Ω. Immediately following electroporation, the cells were diluted in 1 ml of pre-reduced and pre-warmed TSPC. The culture was plated on TSPC agar plates with 5 g/ml thiamphenicol after being incubated anaerobically for 5 hours without agitation.

### Gene deletion based on *hicA (3A*)-counterselecitonin *F. nucleatum*

To create deletion constructs for *radD* that eliminate the amino acids in positions between 44 and 1240, 1.6-kb fragments upstream and downstream of the *radD* interested region were amplified by PCR using primer pairs RadDupF/R and RadDdnF/R (Table 2). For the *tnaA* deletion plasmid, 1.6-kb fragments upstream and downstream of *tnaA* were PCR amplified using primer pairs tnaAupF/R and tnaAdnF/R. An overlapping PCR reaction fused the upstream and downstream fragments. Each fused segment was then mixed in 10 μl Gibson assembly master mix solution with an equal amount of the pBCG02 vector backbone in which the *repA* and *ori*_Fn_ element were removed by inverse PCR with primer pair pCWU6-F/R (see Figure 3A). A 2x hot-start PCR master mix from Takara (cat#: R405A) was used for high-fidelity PCR amplification of the pBCG02 backbone. The resulting plasmids pBCG02-Δ*radD* and pBCG02-Δ*tnaA* were further confirmed by DNA sanger sequencing and transformed into *F. nucleatum* ATCC 23726 by electroporation (52), respectively.

Plasmid insertion into the bacterial chromosome by homologous recombination was selected by growth with TSPC liquid media in the presence of 5 μg/ml thiamphenicol at 37°C. A 5 μg/ml thiamphenicol-resistant colony was inoculated overnight in TSPC without antibiotics. The following day, a 100 μl aliquot of 1,000-fold diluted cultures was plated on TSPC agar plates containing 2 mM inducer theophylline to select for cells in which the plasmid had been excised and lost. Growing 8~10 colonies restreaked on TSPC plates, checked for thiamphenicol sensitivity (100% efficiency), and screened by PCR amplification for the presence of the expected deletion in *radD* or *tnaA*.

### Bacterial co-aggregation

Co-aggregation assays were performed with wild-type, *ΔradD* and 8 colonies from *hicA(3A)-*based counterselection of *F. nucleatum* and *A. oris* MG-1 as previously described (17). Briefly, stationary-phase cultures of bacterial strains were grown in TSPC or heart infusion broth (for MG-1) harvested by centrifugation, washed in coaggregation buffer (53), and suspended to an equal cell density of approximately 2 x 10^9^ ml^-1^ based upon OD_600_ values. For co-aggregation, 0.20 ml aliquots of *Actinomyces* and fusobacterial cell suspensions were mixed in a 24-well plate for a few minutes on a rotator shaker, and images were taken.

### Indole assay

Fusobacterial wild-type and ten clones, obtained through *hicA(3A)-*counterselection targeted towards in-frame deletion of the *tnaA* gene, were individually inoculated into a test tube containing 8 ml of TSPC medium. The tubes were incubated at 37°C for 24 hours, and a 200 μl aliquot was taken from each culture and added to a 96-well plate. Following this, 100 μl of Kovacs’s reagent was added to each tube without shaking, and resulting color change was observed. A positive indole test is indicted by the formation of a pink to red color in the reagent layer on top of the medium within a few seconds of adding the reagent, while a negative result appears yellow. The photograph was taken 10 minutes after the reagent was added.

## AUTHOR CONTRIBUTIONS

C.W. conceived and designed all experiments. B.G. and P.Z. performed all experiments. B.G., P.Z., and C. W. analyzed data. C.W. wrote the manuscript with contribution and approval from all authors.

## DATA AVAILABILITY STATEMENT

Materials are available upon reasonable request with a material transfer agreement with UTHealth for bacterial strains or plasmids. pBCG01 plasmid sequence and map can be accessed on a Benchling link at https://benchling.com/s/seq-VuTCpImziZSBrxT6otvO?m=slm-jUtGU7tIsA1ZnPAgUH1f

## ACKNWLEDGMENTS

This work was supported by a National Institute of Dental and Craniofacial Research (NIDCR) grant (DE030895) to C.W.

